# How Accurate Are Circular Dichroism Based Secondary Structure Estimates?

**DOI:** 10.1101/2020.06.05.123398

**Authors:** Gabor Nagy, Helmut Grubmüller

## Abstract

Circular dichroism (CD) spectroscopy is highly sensitive to the secondary structure (SS) composition of proteins. Several methods exist to either estimate the SS composition of a protein or to validate existing structural models using its CD spectrum. The accuracy and precision of these methods depend on the quality of both the measured CD spectrum and the used reference structure. Using a large reference protein set with high quality CD spectra and synthetic data derived from this set, we quantified deviations from both ideal spectra and reference structures due to experimental limitations. We also determined the impact of these deviations on SS estimation, CD prediction, and SS validation methods of the SESCA analysis package. With regard to the CD spectra, our results suggest intensity scaling errors and non-SS contributions as the main causes of inaccuracies. These factors also can lead to overestimated model errors during validation. The errors of the used reference structures combine non-additively with errors caused by the CD spectrum, which increases the uncertainty of model validation. We have further shown that the effects of scaling errors in the CD spectrum can be nearly eliminated by appropriate re-scaling, and that the accuracy of model validation methods can be improved by accounting for typical non-SS contributions. These improvements have now been implemented within the SESCA package.

## 1 Introduction

Circular dichroism (CD) spectroscopy is known for its high sensitivity to the secondary structure (SS) composition of proteins, especially when bright, synchrotron radiation (SR) light sources are used [5]. CD spectra are routinely used to estimate protein SS compositions, both as a laboratory quality control and to monitor structural changes in proteins. The latter requires the validation of proposed structural models, either by estimating SS compositions from the measured spectra and comparing them to the SS composition of the structural models, or by predicting CD spectra from the structural models and then comparing those to the measured spectra.

Many established SS estimation and CD prediction methods decompose the CD signals into linear combinations of empirical “basis spectra”, representing contributions from SS elements of the protein. The accuracy of these methods depends on several assumptions [2] concerning both measurements of the reference proteins which the basis spectra are extracted from, as well as the measurements on the proteins of interest:

1. *The protein concentrations during CD measurements are accurately known.* To extract accurate basis spectra from different measurements and proteins, the CD spectra need to be properly normalized, which requires an accurate determination of the respective protein concentrations. Unfortunately, the relevant measurements suffer from a 10-25% [8] uncertainty, introducing scaling errors to the measured CD spectra. The propagation of these errors reduces the accuracy of CD prediction and SS estimation methods. Therefore, many methods apply intensity scaling factors to correct the strength of measured CD signals.
2. *The SS composition of reference proteins is accurately known, and reflects the SS composition under the the conditions of the CD measurement.* Methods that rely on empirical basis spectra require reference SS compositions, usually obtained from structural models determined by X-ray crystallography or nuclear magnetic resonance (NMR) measurements. Structure determination typically requires conditions different from those of CD measurements (e.g. different concentrations), which may alter the protein structure. As a result reference SS compositions typically deviate from those of the solution structure by 10 % on average [6], reducing the accuracy of empirical SS estimation and CD prediction methods.
3. *The measured protein samples are free of contamination, and non-SS CD contributions can be neglected.* Non-SS contributions from the protein include tertiary structure CD contributions, far ultra-violet (UV) CD signals from natural or modified amino acid side chains, co-factors, and ion coordination sites. The most studied of those are side chain contributions, which, however are typically smaller than 10% of the SS contributions [12].

Here, we will address two questions: First, to which extent are the above assumptions violated in typical SR-CD data sets? Second, how do such deviations affect the accuracy of SS estimation, CD prediction, and model validation methods? To answer the second question, we constructed a synthetic reference data set including the typical violations, for which the deviations in the reference data are precisely known, and their effects are exactly calculable.

## 2 Methods

### 2.1 Experimental errors

Typical deviations from the assumptions listed above were estimated based on the analysis performed by Nagy *et al.* [12] on the SP175 reference set [7], which contains high-quality structures and SR-CD spectra for 71 proteins with diverse SS compositions.

Briefly, the correct SS composition for the protein in solution was estimated through deconvolution of its re-scaled CD spectrum. The scaling factors applied to the measured spectra quantified scaling errors in the data set. Deviations between the estimated correct SS and the reference SS composition were used to quantify structural errors. Finally, non-SS contributions were quantified by averaging the deviations between the re-scaled CD spectra and CD signals back-calculated from the estimated SS.

The scaling factor *α_j_* for each reference protein was determined based on six predicted spectra, each calculated from the same reference structure using different prediction methods. Four of these predictions were made by SESCA basis sets (DS-dT, DSSP-1, HBSS-3, DS5-4), one was determined by the *ab initio* predictor DichroCalc [1], and one by a specialized basis set BestSel der ref [12]. For each prediction, a scaling factor was calculated to minimize root mean squared deviation (RMSD) between the measured and predicted CD spectrum. The final *α_j_* for the protein *j* was calculated as the average of its six obtained scaling factors, whereas the scaling error of its CD spectrum is given by

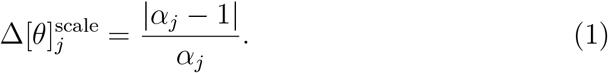

After all reference CD spectra were re-scaled by the *α_j_* values, the four SESCA basis sets were used to obtain the estimated SS composition 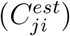 through CD deconvolution. The deviation (Δ*SS_j_*) between the estimated and reference SS compositions were computed according to

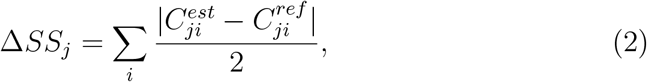

where 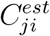 and 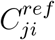 are the coefficients of SS class *i* in protein *j* for the estimated and reference structures, respectively. The obtained Δ*SS_j_* values from each basis set were again averaged for every protein *j* to estimate the SS deviation of reference structures in the SP175 set.

For each protein the estimated prediction error caused by non-SS CD contributions 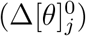 was calculated and normalized by the re-scaled average spectrum intensity

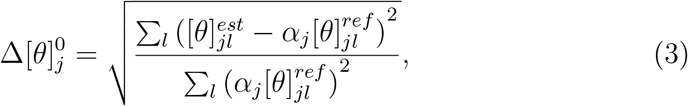

where 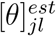 and 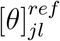 are back-calculated and measured spectral intensities of protein *j* at wavelength *l*, respectively. Similarly to SS deviations, 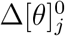 values calculated using the 4 SESCA basis sets were averaged for each protein in the SP175 set to obtain a final estimate on its non-SS contributions.

Next, the noise-to-signal ratio 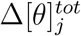 for each reference protein was determined by dividing the total prediction error by the average intensity of the estimated SS signal

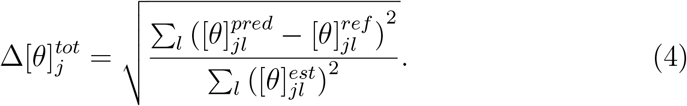

Again, the four obtained values from SESCA basis sets were averaged for each protein *j* to estimate the final noise-to-signal ratio for all reference proteins. The distribution of scaling factors (*α_j_*), SS deviations (Δ*SS_j_*), non-SS contributions 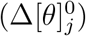, and noise-to-signal ratios 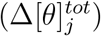 of the SP175 set were used to describe the typical deviations from the assumed ideal experimental data, as well as to generate synthetic data sets that test the effect of these deviations during SR-CD based model validation.

### 2.2 Synthetic Data

A synthetic data set of structures and CD spectra with precisely known errors were created to test the effect of different deviations from the ideal experimental data on the CD prediction, SS estimation, and model validation methods.

A “correct model” was defined with a typical SS composition of 30% *α*-helix, 40% *β*-strand and 30% random coil.

From that model a “correct CD signal” (purple-dashed curve in Fig. 1) was generated by predicting the CD spectrum of the correct model with the DS5-4 basis set of SESCA [12]. For the CD prediction, SS fractions of the correct model were assigned to the coefficients of basis spectrum “Helix1”, “Beta1”, and “Other”, respectively.

**Figure 1:**
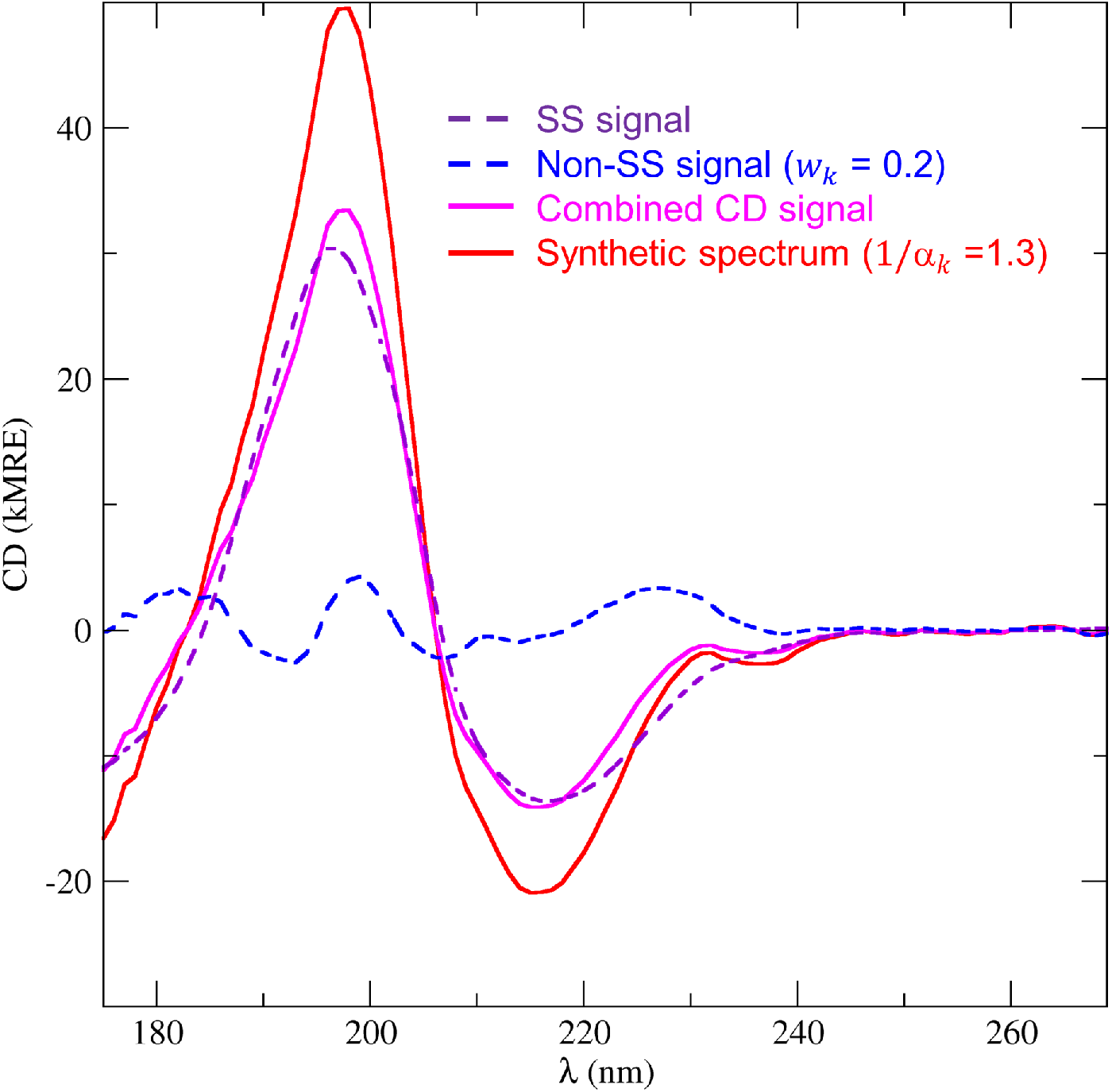
Constructing synthetic CD spectra. Synthetic spectra are constructed from a SS signal (purple-dashed line), a weighed non-SS signal (blue-dashed line), and a scaling factor (1/*α_k_*). The non-SS signal is scaled to a given fraction (*w_k_*, here 0.2) of the average SS signal intensity, then added to the SS signal, to imitate non-SS contributions of different sizes. Finally, this combined CD signal (in magenta) is multiplied by a scaling factor (here 1.3) to mimic scaling errors, yielding the final synthetic spectrum (in red). The weighs and scaling factors for all used synthetic spectra are provided in Table 2.

Structural deviations were modelled by constructing 20 synthetic models with altered SS compositions that covered the *α* – *β* – coil SS space (see Table 1).

**Table 1:** Synthetic models with diverse SS compositions used for error assessment. The table provides the name and the identifier *j* of the model, the fraction of residues classified as *α*-helix, *β*-strand, and other secondary structure classes, as well as the true SS deviation (Δ*SS_j_*) from the correct model (*j*=0) of the synthetic data set.

| Model | $j$ | α-helix | β-strand | Other | $\Delta SS_j$ |
| --- | --- | --- | --- | --- | --- |
| correct | 0 | 0.3 | 0.4 | 0.3 | 0% |
| AB+30 | 1 | 0.0 | 0.7 | 0.3 | 30% |
| AB+20 | 2 | 0.1 | 0.6 | 0.3 | 20% |
| AB+10 | 3 | 0.2 | 0.5 | 0.3 | 10% |
| AB-10 | 4 | 0.4 | 0.3 | 0.3 | 10% |
| AB-20 | 5 | 0.5 | 0.2 | 0.3 | 20% |
| AB-30 | 6 | 0.6 | 0.1 | 0.3 | 30% |
| AB-40 | 7 | 0.7 | 0.0 | 0.3 | 40% |
| BC+36 | 8 | 0.3 | 0.04 | 0.66 | 36% |
| BC+26 | 9 | 0.3 | 0.14 | 0.56 | 26% |
| BC+16 | 10 | 0.3 | 0.24 | 0.46 | 16% |
| BC+6 | 11 | 0.3 | 0.34 | 0.36 | 6% |
| BC-6 | 12 | 0.3 | 0.46 | 0.24 | 6% |
| BC-16 | 13 | 0.3 | 0.56 | 0.14 | 16% |
| BC-26 | 14 | 0.3 | 0.66 | 0.04 | 26% |
| AC+23 | 15 | 0.07 | 0.4 | 0.53 | 23% |
| AC+13 | 16 | 0.17 | 0.4 | 0.43 | 13% |
| AC+3 | 17 | 0.27 | 0.4 | 0.33 | 3% |
| AC-3 | 18 | 0.33 | 0.4 | 0.27 | 3% |
| AC-13 | 19 | 0.43 | 0.4 | 0.17 | 13% |
| AC-23 | 20 | 0.53 | 0.4 | 0.07 | 23% |

**Table 2:**
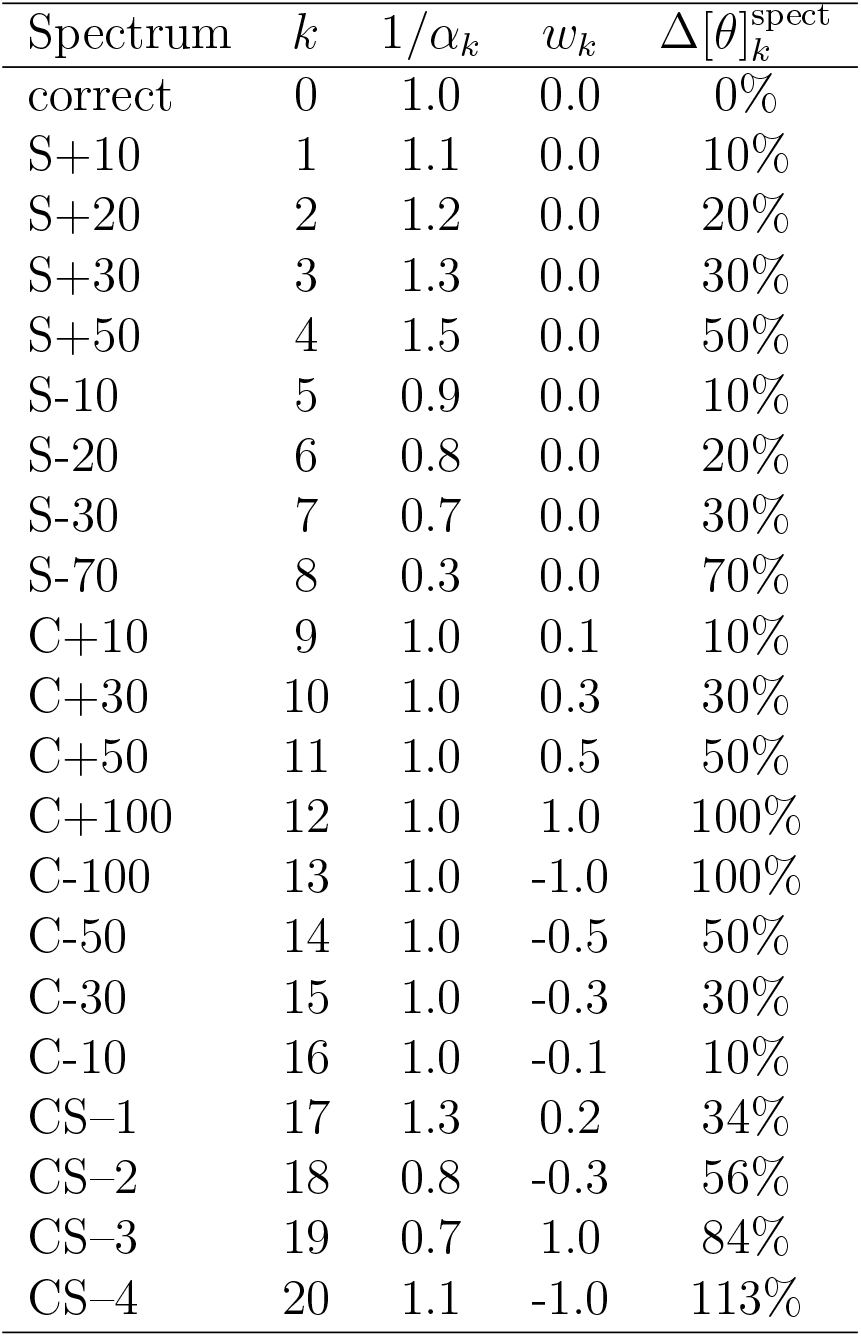
Synthetic CD spectra with diverse CD deviations used for error assessment. The table lists the name and identifier *k* of the synthetic spectra, the scaling factors 1/*α_k_*, and weights *w_k_* used to add scaling errors and non-SS contamination to the correct spectrum (k=0), as well as true deviation 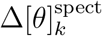 from the correct spectrum (k= 0), expressed as a percentage of the true spectrum intensity.

| Spectrum | $k$ | $1/\alpha_k$ | $w_k$ | $\Delta[\theta]_k^{\text{spect}}$ |
| --- | --- | --- | --- | --- |
| correct | 0 | 1.0 | 0.0 | 0% |
| S+10 | 1 | 1.1 | 0.0 | 10% |
| S+20 | 2 | 1.2 | 0.0 | 20% |
| S+30 | 3 | 1.3 | 0.0 | 30% |
| S+50 | 4 | 1.5 | 0.0 | 50% |
| S-10 | 5 | 0.9 | 0.0 | 10% |
| S-20 | 6 | 0.8 | 0.0 | 20% |
| S-30 | 7 | 0.7 | 0.0 | 30% |
| S-70 | 8 | 0.3 | 0.0 | 70% |
| C+10 | 9 | 1.0 | 0.1 | 10% |
| C+30 | 10 | 1.0 | 0.3 | 30% |
| C+50 | 11 | 1.0 | 0.5 | 50% |
| C+100 | 12 | 1.0 | 1.0 | 100% |
| C-100 | 13 | 1.0 | -1.0 | 100% |
| C-50 | 14 | 1.0 | -0.5 | 50% |
| C-30 | 15 | 1.0 | -0.3 | 30% |
| C-10 | 16 | 1.0 | -0.1 | 10% |
| CS-1 | 17 | 1.3 | 0.2 | 34% |
| CS-2 | 18 | 0.8 | -0.3 | 56% |
| CS-3 | 19 | 0.7 | 1.0 | 84% |
| CS-4 | 20 | 1.1 | -1.0 | 113% |

CD deviations were modelled by constructing 20 synthetic CD spectra with scaling errors, non-SS contributions or both (Table 2).

Scaling errors were modelled multiplying the correct spectrum with 1/*α_k_* = {0.3, 0.7, 0.8, 0.9, 1.1, 1.2, 1.3, 1.5} to obtain four under-scaled (subsequently S−) and four over-scaled (S+) CD spectra.

Errors from non-SS CD contributions were modelled by adding a “contamination” signal (blue-dashed curve in Fig. 1) to the correct spectrum. The contamination signal was obtained by estimating the SS composition of bovine lactoferrin (SP175/42) from its measured CD spectrum, and subtracting its estimated SS signal from the measured one. This contamination was re-scaled to the same average intensity as the correct spectrum, and then was added to the correct spectrum with weights of *w_k_* = {±0.1, ±0.3, ±0.5, ±1.0} to create two series of CD spectra (C+ and C−) with increasing non-SS contributions.

Further, a set of four CD spectra (CS) was generated that included both contamination and scaling errors. For these spectra, weights *w_k_* = {0.2, −0.3, 1.0, −1.0} were used to add contamination, then the resulting spectra were scaled by 1/*α_k_* = {1.3, 0.8, 0.7, 1.1}, respectively.

The error in each synthetic spectrum *k* was calculated and normalized by the correct CD signal

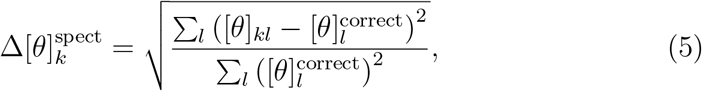

where [*θ*]_*kl*_ and 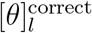 are CD intensities of spectrum k and the correct spectrum at wavelength *l*, respectively.

### 2.3 Deconvolution methods

We used three different deconvolution methods termed D1, D2, and D3 to study the effects of experimental errors on SS estimation accuracy. All three methods use the DS5-4 basis set and perform several simplex searches in the SS composition space based on an adaptive Nelder-Mead algorithm [3], as implemented in the deconvolution module of the SESCA package [12]. The three methods differ in the number of searches performed as well as in the applied constraints as described below. We note that the application of such constraints reportedly affects the accuracy of the deconvolution, depending on the experimental error of the CD spectrum of interest [9].

For D1, 500 simplex searches were performed, each starting from a random SS composition. As constraints, each basis spectrum coefficient was required to be non-negative and their sum to be unity. For D2, the sum of coefficients was not required to be unity and, due to faster convergence, only 200 searches per protein were performed. D3 proceeds as D2, except the coefficients are not restricted to non-negative values during the search.

At the end of the deconvolution, the search resulting in basis set coefficients with the best fit to the measured spectrum was accepted. For the accepted fit, all negative coefficients were set to zero, and subsequently coefficients were re-normalized to add up to unity. This procedure yielded plausible SS compositions for all three methods, and also provided the optimal scaling factors for the measured spectra for D2 and D3.

### 2.4 Model validation methods

We tested the accuracy five potential validation methods, which may be used to evaluate the quality of protein structural models with SESCA [12]. Three methods (V1, V2, and V3) are based mainly on CD deconvolution, the other two (V4 and V5) are based on CD predictions.

Specifically, V1 estimates the SS composition of a target protein without corrections to the CD spectrum, using deconvolution method D1. The error of its proposed model 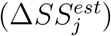 is then calculated according to eq. 2. Method V2 is similar to V1, except that the deconvolution is done by D2, which includes re-scaling the measured CD spectrum during the SS estimation. Method V3 is also similar to V1, except that prior the deconvolution step, the measured CD spectrum is re-scaled to match the intensity of the predicted CD spectrum of the proposed model. We note that method D3 was not considered for model validation based on its sensitivity to non-SS contributions discussed in Section 3.2.

V4 and V5 first predict CD spectra from the proposed protein structure, then calculate 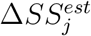 from the deviation of the predicted and measured CD spectra (*RMSD_j_*) according to

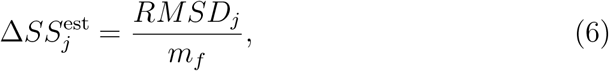

where *m_f_* is a predetermined sensitivity parameter. For both methods, the measured CD spectrum is re-scaled to minimize the *RMSD_j_* prior the estimation of the model error. The two methods differ in their sensitivity parameters, which was *m_f_* = 15.6 kMRE for V4 and *m_f_* = 30.7 kMRE for V5. The former was determined based on a calibration using the SP175 set as described in [12], whereas the latter was derived using the same calibration performed on a set of 500 random generated synthetic reference proteins, that mimicked the distribution of SS compositions, estimated scaling errors and non-SS contributions of the SP175 set (the latter two distributions are discussed in Section 3.1).

### 2.5 Model validation accuracy

The accuracy of all model validation methods described above was evaluated from the synthetic data set described in Section 2.2 using two different metrics. First, the model validation error for a given synthetic CD spectrum *k* was calculated as

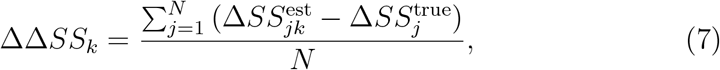

where 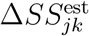 is the estimated SS deviation between model *j* and the correct model, determined using spectrum *k*, and 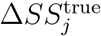 is true SS deviation listed in Table 1. Second, a ranking score *R_k_* was determined, which quantifies how many of the other 20 synthetic models had 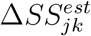 values lower or equal to the correct model. Both ΔΔ*SS_k_* and *R_k_* values were computed systematically for each CD spectrum in the synthetic data set, to assess the change in model validation accuracy as function of experimental errors in the reference CD spectrum. Finally, the mean and standard deviation of model errors (ΔΔ*SS*) and ranking scores (Avg. Rank) were computed to quantify the overall performance of the method.

## 3 Results

### 3.1 Experimental error distribution

First, we characterized the typical deviations from the three assumptions that define ideal SR-CD data (Section 1). These deviations were quantified for 71 reference proteins of the SP175 set [7] through scaling errors and non-SS contributions of their measured CD spectra (collectively referred to as CD deviations), as well as through SS deviations between their structural model and estimated correct structure (Section 2.1).

Figure 2A shows the distribution of the scaling factors *α_j_*, i.e. the ratios of assumed and correct protein concentrations, that compensate for estimated scaling errors in the SR-CD spectra. As expected from random errors due to measurement uncertainty, the distribution of the *α_j_* values is close to normal, with a mean of 0.87 and a standard deviation (SD) of 0.25. The SD also agrees well with the typical uncertainty reported for protein concentration measurements [8, 4]. The fact that the mean value is smaller than unity is likely due to protein adsorption at the cell surface during the CD measurements, effectively decreasing the actual concentration in the bulk.

**Figure 2:**
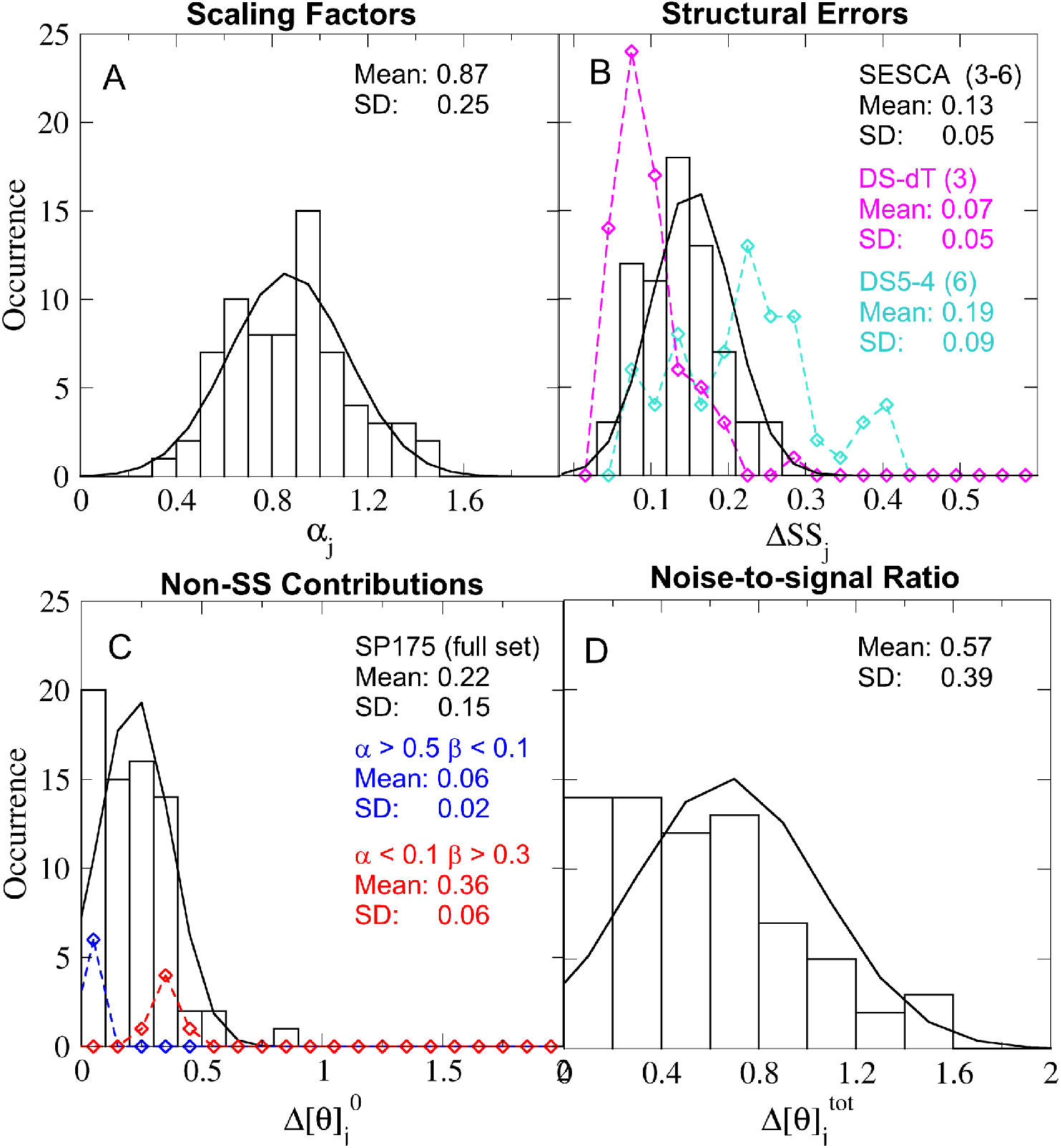
Estimated error distribution in the SP175 reference set. Histograms show the binned distribution of estimated intensity scaling factors due to incorrect normalization (A) and non-SS contributions (C) of the measured CD spectra in the reference set, as well as the fraction of miss-classified amino acids in the reference structures (B) and noise-to-signal ratios (D) of the predicted CD spectra caused by the above three factors. Solid lines indicate expected occurrences assuming Gaussian fits. The turquoise and magenta symbols in panel B show the distribution of SS deviations estimated using only the DS-dT and DS5-4 basis sets with three and six basis spectra, respectively (the black distribution is an average over four basis sets). The blue and red symbols in panel C show non-SS contributions for the *α*-helical and *β* + coil sub-populations of SP175, respectively.

Figure 2B shows the SS deviations Δ*SS_j_*, calculated as an average over predictions from four different SESCA basis sets (Section 2.1). These are also close to be normally distributed, with a mean of 0.14 and a SD of 0.05. These SS deviations between the reference structures and the SS composition derived from the measured CD spectra are larger than the 10% expected from comparing X-ray structures and NMR structures of the the same protein [9]. Note, however, that expected 10% deviation is based on a classification of only three SS classes, whereas the four SESCA basis sets have three to six SS classes. The mean SS deviation over all reference proteins computed for individual basis sets increases monotonically with the number of SS classes from 7% for three SS classes (magenta) to 19% for six SS classes(cyan), which may explain the obtained larger average deviations. However, we also note that the uncertainty of the estimated correct SS compositions derived from the CD spectra (Section 2.1) may also contribute to the obtained SS deviations.

Figure 2C shows the distribution of non-SS contributions 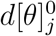, estimated from the difference between the SS contribution derived from deconvolution and the (re-scaled) measured spectrum (see Section 2.1). Clearly, the Gaussian fit (black line) does not describe this distribution well. For about half of the reference CD spectra, the non-SS contributions are smaller than 20% of the CD signal intensity, consistent with the assumption that, for these cases, the signal is dominated by the SS contributions. However, for the rest of the proteins, larger non-SS contributions of up to 60% are seen, with one outlier close to 80%. We note that non-SS contributions tend to be smaller for *α*-helical proteins (blue symbols) than for *β*-sheet and Coil proteins, due to the stronger CD signal of *α*-helices. Further, due to the fitting procedure used to estimate the correct SS compositions, the histogram in Figure 2C rather underestimates the actual deviations. These findings render the question of how the non-SS contributions affect the interpretation of CD spectra particularly relevant. We will address this question further below.

To quantify the combined effects of the above three deviations, the noise-to-signal ratios 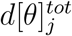 were also calculated for each reference protein. These ratios, similarly to the non-SS contributions, are not normally distributed, and a wide range of ratios between 0.1 to 1.6 was obtained for the SP175 set. This distribution also shows that, even with the best experimental information available, the noise caused by non-ideal experimental data is larger than 40% of the SS signal for over half of the studied reference proteins.

Considering the estimated noise levels, it is surprising that in our previous study [12] the accuracy of SESCA basis sets appeared to be robust to errors in the SP175 reference set. This robustness is likely observed because the basis spectra are determined from a large set of structures and CD spectra, and the influence of errors from individual proteins is largely reduced due to averaging. However, during model validation, we cannot rely on such cancellation of errors in the reference CD spectrum and the SS composition of the protein of interest (henceforth, target protein). Therefore, the remaining sections will focus on the effect of CD and SS deviations of the target protein with respect to the accuracy of SS estimation, CD prediction, and model validation methods.

### 3.2 Effects on the accuracy of SS estimation methods

First, we tested how CD and SS deviations affect the accuracy of the three SS estimation methods D1, D2, and D3, described in Section 2.3. All three methods estimate the SS composition of the target protein by spectrum deconvolution, approximating its measured CD spectrum with a linear combination of basis spectra. The methods differ in the constraints applied to the basis spectrum coefficients during the search for the best approximation. D1 applies both normalization and non-negativity constraints to the coefficients, D2 only applies the non-negativity constraint, and D3 applies no constraints.

As a first step, we consider the effects of CD deviations on the accuracy of SS estimation methods, because these deviations directly affect CD deconvolution. Then, as a second step, we illustrate how the errors from CD deviations and SS deviations in reference structures combine for model validation methods based on SS estimation, such as the scheme we used to estimate CD and SS deviations in Section 3.1.

We test the effect of CD deviations in the target spectrum by gauging the accuracy of SS estimates from 21 synthetic CD spectra, to which we intentionally introduced given amounts of scaling errors and non-SS contributions (listed in Table 2). The error of methods D1, D2, and D3 for synthetic (target) spectrum *k* was determined from the deviation Δ*SS_k_* between their SS estimate and the correct SS composition of the synthetic data set. Figure 3 shows how these errors increase in response to different CD deviations. Comparing the errors we obtained from synthetic spectra with the same type of CD deviations (corresponding colours and symbols) highlights that the SS estimation accuracy strongly depends on the applied constraints as well as the CD deviation type.

**Figure 3:**
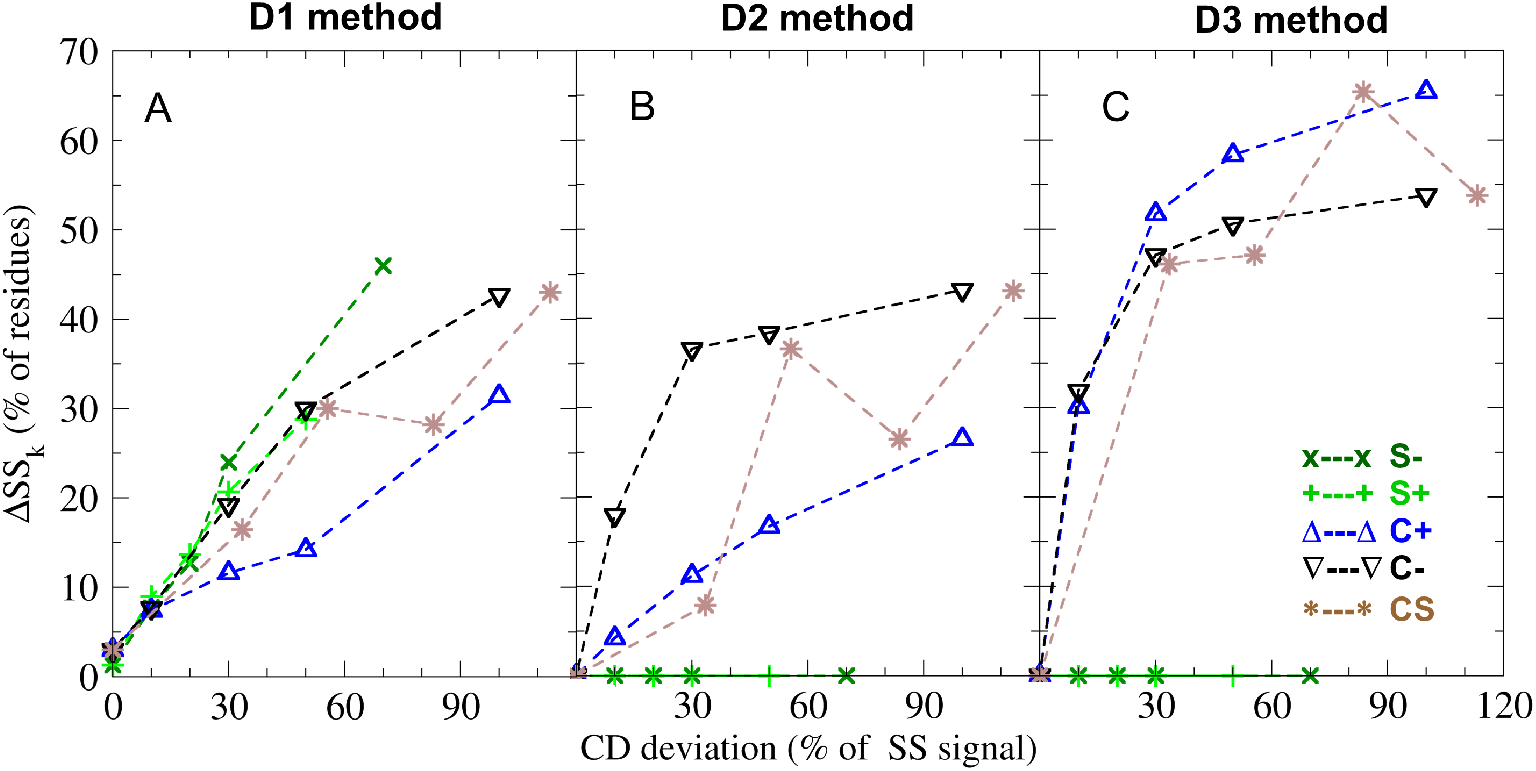
Accuracy of SS estimation methods. The panels show deviations between the estimated and correct secondary structure composition (Δ*SS_k_*) as a function of errors in the reference CD spectrum for deconvolution methods D1 (A), D2 (B), and D3 (C), described in Section 2.3. Light green and dark green symbols denote under-scaled (S−) and over-scaled (S+) CD spectra, blue and black triangles depict CD spectra with two types of non-SS contamination signals (C+ and C−), and brown star symbols denote spectra with both scaling and contamination errors (CS), respectively (see Section 2.2). The error of the spectrum is expressed as a percentage of the correct secondary structure signal.

We first focus on the errors of method D1 (Fig. 3A). This method constrains the basis set coefficients to be positive and sum up to unity, such that the coefficients are equal to the fraction of amino acids in a particular SS class. The obtained Δ*SS_k_* for D1 average to 20.4% and increase almost linearly up to a 25% deviation in the target spectrum. At larger CD deviations, D1 shows a slightly higher sensitivity to scaling errors (S+ and S-subsets shown in light and dark green) than to non-SS contamination (C+ and C−, in blue and black). For synthetic spectra with both scaling errors and non-SS contributions (CS, in brown), the SS estimation error for D1 remains moderately large and changes nearly linearly with summed CD deviation. We note that, despite its limited accuracy, several methods (including SESCA) enforce similar constraints as D1 during their SS estimation. Further, the D1 SS search did not always converge due to the applied constraints. Because the search minimizes the error of the approximation, the error values obtained for D1 are likely overestimated. Non-convergence is also the likely reason for the 1.3% SS deviation observed at 0% spectrum error. Figure 3B shows the same analysis for method D2, which only applies non-negativity constraints, and re-normalizes the best fitting coefficients at the end of the SS search. Because this procedure effectively re-scales the measured CD spectrum during the search, it eliminates SS estimation errors from scaling errors. However, as seen from the errors of the C+ and C− subsets, D2 shows an increased sensitivity to non-SS contamination. The considerable difference of Δ*SS_k_* obtained for the C+ and C− spectrum subsets also indicates that D2 is more sensitive to the shape of the contamination signal. Overall, D2 still yields the smallest average error of 14.4% for the synthetic data set. The better accuracy may explain why some of the more recent deconvolution algorithms (e.g., BestSel [11]) are based on similar constraints.

Carrying the idea of relaxing constraints one step further, it has been suggested [9] not to constrain the coefficients at all during the spectrum approximation (method D3, Fig. 3C). The errors obtained for D3 are zero for S+ and S− subsets, but larger than 30% for all other synthetic spectra, leading to an average SS estimation error of 27.3%. These Δ*SS_k_* values indicate that D3 also eliminates the effect of scaling errors, but it is much more susceptible to over-fitting due to non-SS contributions. Based on these large SS estimation errors and the distribution of non-SS contributions reported in Section 3.1; we expect method D3 to be rather inaccurate for about one third of the proteins of the SP175 set. Consequently, we decided not to analyse methods using unconstrained deconvolution further.

The obtained results enable us to determine how much the estimated SS compositions, on average, differ from the true solution structure of the CD measurement as a function of CD deviations in the spectrum. The synthetic data set also allows us to assess how these differences affect model validation methods based on SS estimation. To this aim, we consider the combined effect of errors in the SS estimation and the error in the structural model(s) to be validated (e.g. reference structures from X-ray crystallography). To test the combined effect of these errors, in a second step, we use 20 synthetic SS compositions with different SS deviations from the correct structure (see Table 1). In Figure 4, these synthetic models play the role of experimental ‘known’ structures that are compared to the estimated SS composition based on the CD spectrum.

**Figure 4:**
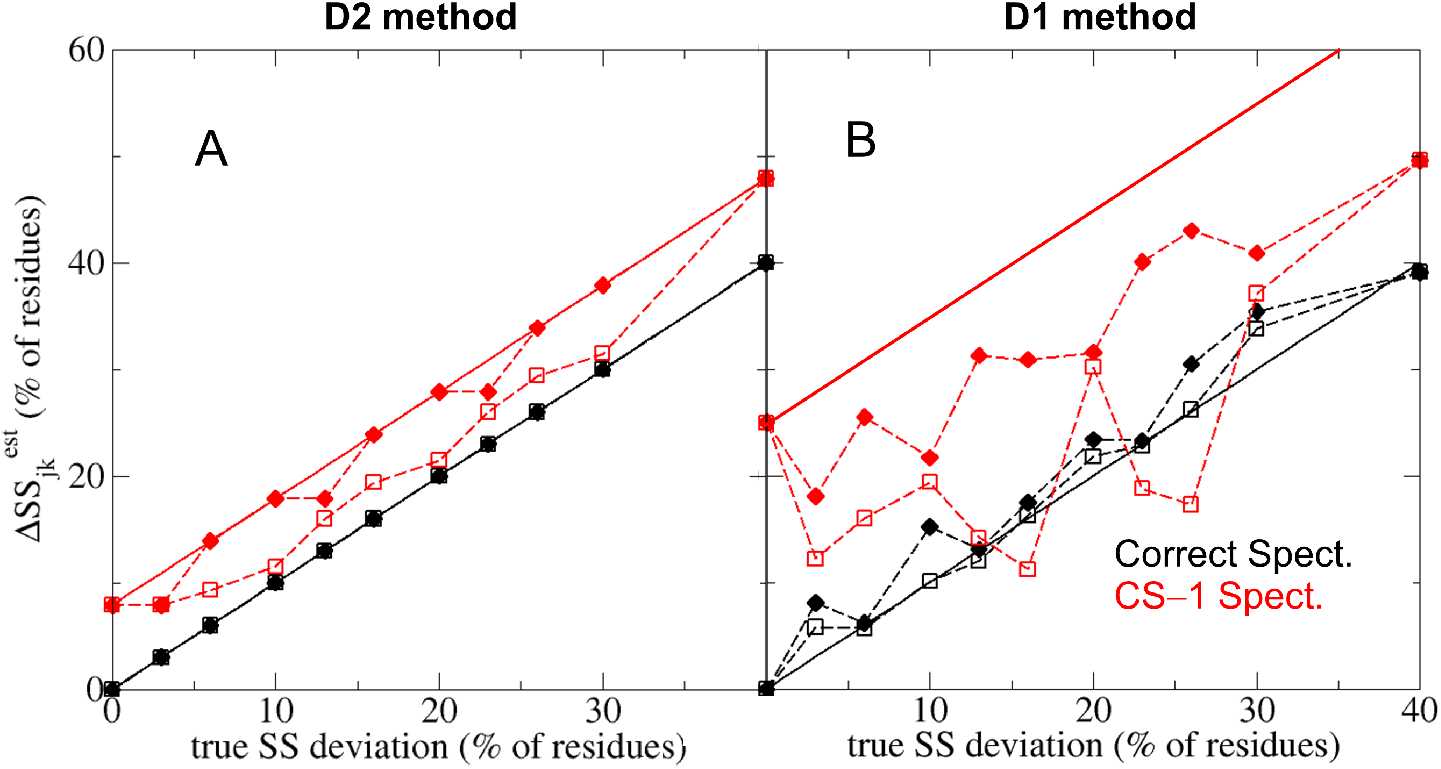
Errors in the reference structure affect model validation. The estimated SS deviation (Δ*SS_jk_*) between the reference and correct structure is shown as function of the true SS deviation for deconvolution methods D2 (A) and D1 (B). The symbols depict the smallest (empty squares) and largest (full diamonds) estimated SS deviations for synthetic reference models of a given true SS deviation. Symbols in black denote SS estimates based on the correct CD spectrum of the set (no CD deviations), whereas red symbols were based on a spectrum with typical CD deviations (CS–1, see Table 3). The black solid lines show the expected SS deviation based on accurate SS estimates. Red solid lines indicate expected estimated SS deviations, if the errors caused by the CD and SS deviations were additive.

Initially, we estimate the true SS composition from the correct synthetic CD spectrum (k=0 in Table 2, no scaling errors or non-SS contributions) to determine 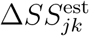, the estimated SS deviation of each synthetic model *j*. In Figure 4A and 4B, these estimated SS deviations (black symbols) are shown for methods D2 and D1, respectively, as function of the true SS deviation. Because D2 always estimates the true SS composition accurately from the correct synthetic spectrum, the estimated model errors are equal to the true SS deviation, and the black symbols in Fig. 4A fall on the black solid line that indicates an accurate model validation. The estimated SS deviations for D1 (Fig. 4B) differ slightly from the true SS deviations on several instances, most likely because the SS estimation does not always converge. Overall, for using an ideal CD spectrum, the correct SS compositions are exactly or almost exactly recovered by the two methods and, therefore, in this case, the observed SS deviations only – and trivially – reflect the difference between the reference and true SS compositions.

**Table 3:** Average performance of validation methods. The table lists the name of the method, its average model validation error (ΔΔ*SS*) and standard deviation (SD), as well as the average and SD of its ranking score (Avg. Rank).

| Method | $ddSS$ (%) | SD (%) | Avg. Rank | SD |
| --- | --- | --- | --- | --- |
| V1 | 14.4 | 8.4 | 4.9 | 3.4 |
| V2 | 11.5 | 4.9 | 2.8 | 2.3 |
| V3 | 11.7 | 9.2 | 3.0 | 3.3 |
| V4 | 24.5 | 15.0 | 2.9 | 3.1 |
| V5 | 3.3 | 7.3 | 2.9 | 3.1 |

Next, we estimate the SS deviations from the true structure using a synthetic CD spectrum with CD deviations typical for the SP175 set (CS–1, k=17 in Table 2), which cause a 7.9% and 25% error in the estimated SS composition of D2 and D1 respectively. We attribute this large difference in the SS estimation error to the fact that D2 compensates for the 30% scaling error in the spectrum, whereas D1 does not. If we expect the errors from CD and SS deviation to be additive, then estimated SS deviations should fall on the solid red lines, for which the offsets are errors of the SS estimation. However, the obtained 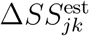 values (red symbols) indicate that the effect of CD and SS deviations are often not fully additive, because the estimated SS deviations are usually larger than the true SS deviation, but by less than the SS estimation error. These results suggest that CD deviations generally lead to an overestimation of the true SS deviation, which increases with the error of the applied SS estimation method.

Further, the dashed lines in Fig. 4 connect the smallest (empty symbols) and largest (full symbols) estimated SS deviations in the synthetic data set observed for a given true SS deviation. The difference between the minimum and maximum estimated SS deviations is zero for accurate SS estimations (Fig. 4A black lines) and increases with the SS estimation error up to 26% (Fig. 4B red lines). In addition, some estimated SS deviations in Fig. 4B are even smaller than the true SS deviation indicating a cancellation of errors. The obtained data suggest that the non-additive summation of errors from CD and SS deviations introduces and uncertainty during model validation, which also increases with the error of the SS estimation. Potentially, the estimated SS deviation for any SS composition may change between its true SS deviation plus or minus the SS estimation error. When CD deviations cause large errors in the SS estimate, this uncertainty may mislead the model validation and prevent the precise determination of the correct SS composition. The results also highlight the importance of re-scaling the CD spectra to reduce the uncertainty from scaling errors, and to improve the precision of model validation.

### 3.3 Effects on the accuracy CD predictions

We also tested the effect of SS and CD deviations on the accuracy of CD prediction methods. These methods compute CD spectra from proposed model structures of the target protein, and the predicted spectra can be compared to a measured reference spectrum for model validation. We note that CD prediction methods are affected by errors in the proposed protein models (i.e. the SS deviation between the proposed and correct structure), but CD deviations in the reference spectrum do not influence their predictions directly. However, scaling errors or non-SS contributions cause deviations between the predicted and measured CD spectra and, therefore, they reduce the prediction accuracy and interfere with model validation.

In Figure 5 we show the CD prediction accuracy quantified by two common metrics. First, the root mean squared deviation (*RMSD_j_*) of CD intensities between the compared spectra of protein *j*, and second, a normalized version (*NRMSD_j_*) by Mao *et al.* [10], where the RMSD is divided by the RMS of the measured CD intensities. The RMSD quantifies the absolute deviation between the measured an predicted spectra, whereas the NRMSD is a relative deviation with respect to the measured CD signal.

**Figure 5:**
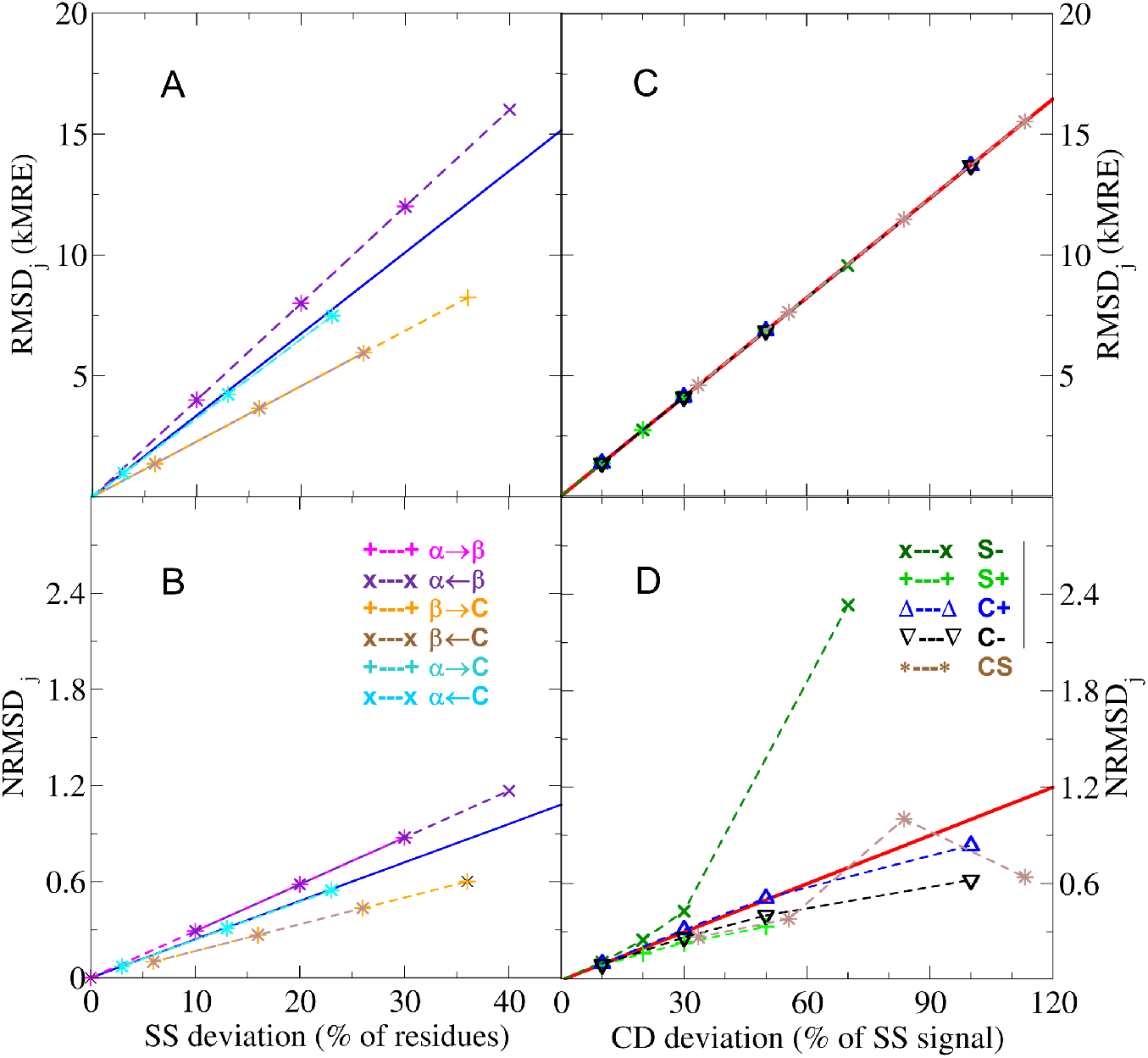
Accuracy of CD spectrum predictions. RMSD (A and C) and NRMSD (B and D) values quantify the accuracy as the deviation of a predicted CD spectrum from a reference spectrum. Panels A and B show the accuracy of CD spectra predicted from synthetic SS compositions with a given error (SS deviation), and compared to the correct reference CD spectrum. Panels C and D show deviations of the CD spectrum predicted from the correct SS composition, compared to reference CD spectra with a given error (CD deviation). The coloured symbols indicate different types of structural and spectral deviations. The symbols in panels A and B denote changes between the fraction of *α*-helices (*α*) to *β*-strands (*β*) and Random coils (C). The symbols in panels C and D denote under-scaled (S-) or over-scaled (S+) CD spectra, spectra with two types of non-SS contamination signals (C+ and C-), and spectra with both scaling and contamination errors (CS). The blues lines in panels A and B show the best linear fit on all SS deviation and RMSD/NRMSD pairs. The red line in panel C shows a linear fit on all CD deviation and RMSD pairs, whereas the red line in panel D indicates the same linear fit with RMSD values normalized by the intensity of the correct CD spectrum.

In Figures 5A and 5B, we depict the effect of SS deviations in the protein model by predicting CD spectra for all 21 SS compositions of our synthetic data set and comparing them to the correct CD spectrum of the set. In the absence of CD deviations, both *RMSD_j_* and *NRMSD_j_* values are linearly correlated with the SS deviation of synthetic model *j*, with a slope that depends on which SS fractions deviate from the correct model. Additionally, the prediction accuracy using both metrics can be approximated from the SS deviation with a single linear function (Pearson correlation coefficient of 0.917), in agreement with the model validation results in our previous study [12].

Figures 5C and 5D show the change in *RMSD* and *NRMSD*, respectively, in response to increasing CD deviations. Here, the CD spectrum was predicted from the correct SS composition and compared to all 21 synthetic CD spectra with given scaling errors and non-SS contributions (see Table 2). The *RMSD_j_* values in Fig. 5C increase linearly with an identical slope for all five subsets of generated CD spectra, indicating that this metric is invariant to the type of the CD deviation. In contrast, the increase of *NRMSD_j_* values in Fig. 5D are non-linear and depend on the error type, because CD deviations affect the normalization term (i.e. the spectrum intensity) differently. Accordingly, the change in *NRMSD_j_* values is superlinear when the measured spectrum intensity is underestimated (S−), but sublinear for spectra with non-SS contributions (C+ and C−) and overestimated spectrum intensities (S+).

We also tested the combined effect of SS and CD deviations through synthetic spectrum and SS model pairs that include both. The observed *RMSD_j_* and *NRMSD_j_* values for these combinations clearly show that the effect of CD and SS deviations are not additive for CD predictions, and introduces a similar uncertainty to the model validation as observed for SS estimation methods in Section 3.2. Despite their non-additivity, the square sum of the errors from CD and SS deviations show a Pearson correlation of 0.953 with the square of total *RMSD_j_*. This behaviour is expected for CD spectra with independent error components, as discussed in our previous study [12]. A similar trend is also observed for *NRMSD_j_* values, with a weaker Pearson correlation (0.911) and a slope smaller than unity (0.88). Because the non-linear response to CD deviations leads to a more complex *NRMSD* profile, we only consider *RMSD*-based prediction methods for the subsequent assessment of the effects on model validation.

### 3.4 Comparison between model validation methods

Finally, we compared the accuracy and reliability of five structural model validation methods (see Section 2.5) with respect to certain deviations in the reference CD spectrum. Our aim is to determine which method is most suitable for assessing the quality of model structures using SESCA [12].

Three of the validation methods (V1-V3) are based on the deconvolution of the validation spectrum, and subsequently computing the difference between the estimated SS composition and the SS of the proposed models. From these methods, V1 and V2 use deconvolution methods D1 and D2 (Section 3.2), respectively, to estimate the correct SS composition. The comparison of V1 and V2 illustrates how re-scaling the CD spectrum intensity affects model validation. Method V3 mimics the model validation scheme we used to estimate typical CD and SS deviations in Section 3.1. This method first re-scales the measured CD spectrum based on the spectrum predicted from the model structure, then estimates the correct SS composition using D1 to compare it with that of the model. The other two methods, V4 and V5, are based on CD predictions. They both re-scale the CD spectrum, and estimate the model error from the deviation between model’s predicted spectrum and the validation spectrum using a sensitivity parameter. The two methods differ in this parameter, which was extracted from experimental reference data for V4, and from synthetic data for V5, respectively (see Section 2.4).

The average performance of each validation method was assessed based on the 441 possible spectrum/model combinations of the synthetic data set. First, for each method, we estimated the model error (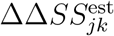, Section 2.5) for every synthetic model *j* based on synthetic spectrum *k*, and compared it to the true SS deviation of the used model from the correct model of the set. The obtained model validation errors were averaged for each spectrum (ΔΔ*SS_k_*) to determine how the CD deviations in the validation spectrum affect the errors of the model validation method. We used the collection of computed ΔΔ*SS_k_* values with increasing CD deviations (henceforth, error profile) to describe the behavior of each method.

Figure 6A shows the error profile of all five methods (dashed lines) for the C-subset of synthetic spectra to illustrate the effect of non-SS contributions in the reference CD spectrum. Overall, the model validation error correlates positively with non-SS signals in the spectrum. The observed increase of ΔΔ*SS_k_* is almost linear for the prediction-based methods (V4 and V5). In contrast, it increases faster at lower errors for deconvolution-based methods (V1-V3), but more slowly at large spectrum errors. In particular, the largest increase is seen for V2, which is not unexpected considering that the underlying D2 deconvolution method shows a larger sensitivity to non-SS contributions.

**Figure 6:**
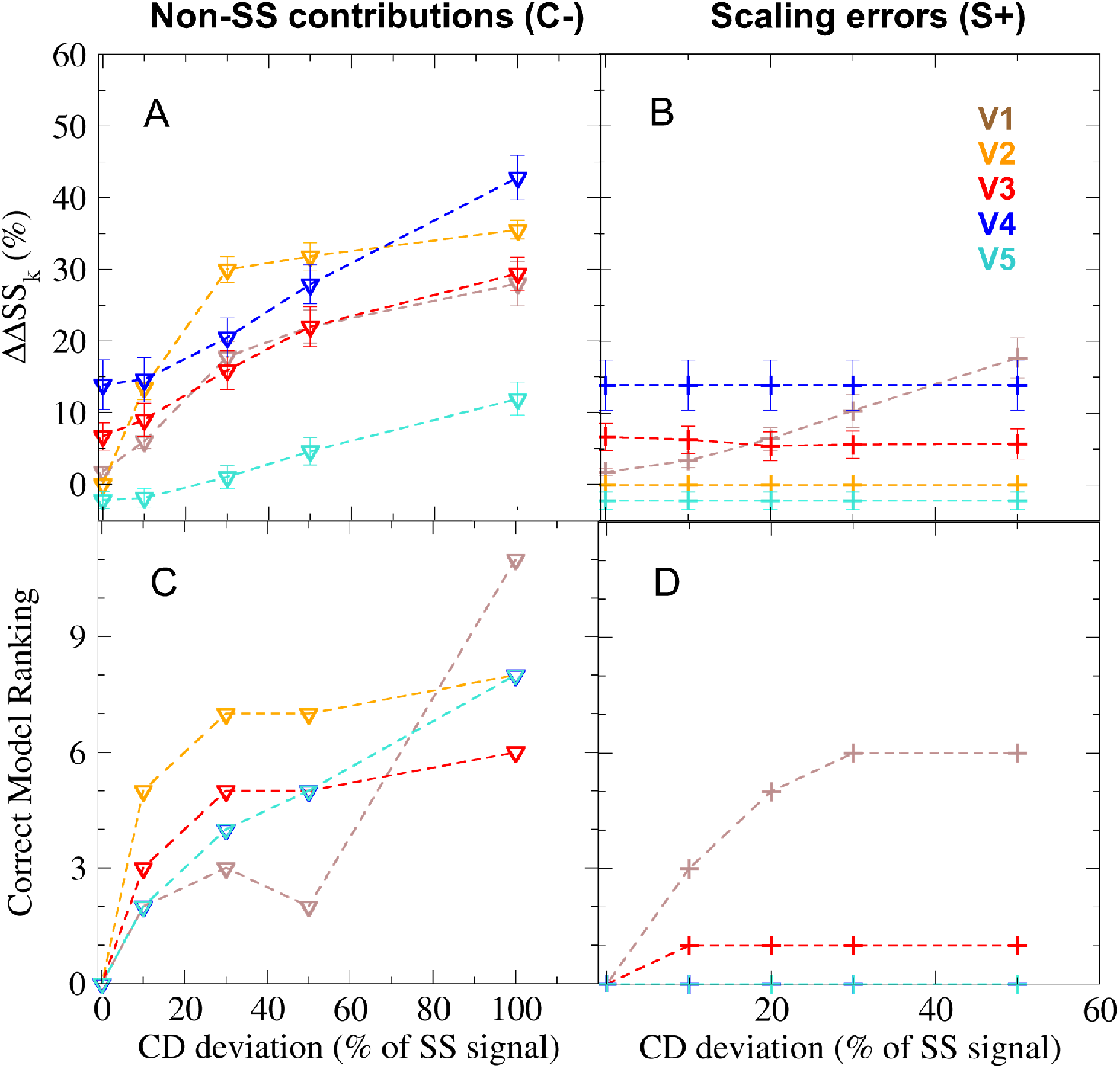
Accuracy and reliability of model validation methods. Validation results shown for the C-(triangles) and S+ (pluses) subsets of synthetic spectra, representing the effects of non-SS contributions and scaling errors, respectively. Results from model validation methods V1 to V5 are depicted in different colours (shown in panel B). The accuracy of validation methods (panels A and B) is quantified by the average difference (ΔΔ*SS_k_*) between the estimated and true errors of the SS composition. These values are computed over 21 synthetic SS models for each synthetic reference CD spectrum and shown as a function of the error in the spectrum (CD deviation). The standard error of ΔΔ*SS_k_* values is shown as error bars. The reliability of the validation methods (panels C and D) is quantified by a ranking score for each reference spectrum, determined by the estimated error of the correct SS model, compared to that of other models. The error in the CD spectrum is expressed as the percentage of the correct secondary structure signal.

It is also informative to analyse the model validation error in the absence of CD deviations (i.e. the offset of the error profiles). Because deconvolution-based model validation always assumes negligible non-SS signals in the CD spectrum, the offset for V1 to V3 is expected to be zero. This is indeed the case for V2, whereas for V1, an average 1.8% deviation is introduced due convergence problems. For V3, re-scaling the CD spectrum to match the predicted spectra of incorrect models introduces an even larger offset of 7%. The prediction-based model validation methods assume an average (non-zero) CD deviation, which should lead to a negative offset in their profiles. This is indeed observed for V5 (−2%), but not for method V4, for which the offset was 14%. The most likely explanation for this large positive offset is an incorrect sensitivity parameter, which for V4 was determined by calibration using estimated SS deviations of the SP175 reference set. We note that these estimated deviations were determined by a modified version of method V3 (see Section 3.2), which, at typical CD deviations (30%) overestimates the model errors by approximately 15%. The propagation of this error to V4 through its sensitivity parameter would explain the observed offset as well as the large difference between the sensitivity parameters of V4 and V5.

We assess the effect of scaling errors in reference CD spectra in Fig. 6B, which shows error profiles for the S+ subset of synthetic spectra. These error profiles have the same offsets but different increase compared to the profiles for non-SS contributions. For V1, which does not re-scale the validation spectrum, model validation errors increase almost linearly with scaling errors, whereas ΔΔ*SS_k_* remains nearly constant for V2 to V5. This trend strongly suggests that re-scaling the reference spectrum indeed eliminates the effects of scaling errors during model validation.

To provide an overall measure of accuracy for the studied validation methods, we also computed the mean and SD of all obtained model validation errors (ΔΔ*SS*, Section 2.5). As Table 3 shows, method V5 predicts the error of synthetic models with the highest accuracy with ΔΔ*SS*= 3.3%, followed by the three deconvolution-based methods V2, V3, and V1 with 11.5%, 11.7%, and 14.4%, respectively, whereas the lowest accuracy is achieved by method V4 (ΔΔ*SS*=24.5%). Note, that the individual model validation errors vary greatly between the model/spectrum pairs for most methods, as shown by their considerable 5-15% standard deviations from the average ΔΔ*SS*. This variation can mainly be attributed to the uncertainty caused by the non-additive summation of errors from CD and SS deviations, which increases with the CD deviations of the validation spectrum.

The presented model validation errors allow us to draw a number of conclusions. First, positive ΔΔ*SS* values indicate that all five methods overestimate the average error of synthetic models. This fact is not unexpected, given that the synthetic data set contains slightly larger than typical CD and SS deviations, due to the over-representation of extreme test cases.

Second, the largest contribution to model validation errors is due to the assumption that CD spectra are solely defined by the SS composition of the protein. Because considerable non-SS contributions are found for more than half of the tested reference proteins, this assumption likely leads to the overestimation of model errors for deconvolution-based methods. Further, the better average accuracy of method V5 indicates that assuming an average non-SS contribution improves model validation significantly.

Third, V4 over-estimates the error of the synthetic models considerably. This result is particularly important, since V4 is the current model validation method of SESCA. This inaccuracy had not been detected so far, because both the calibration and the cross-validation of the method was based on estimated SS deviations using CD deconvolution, which led to a cancellation of errors. This conclusion also suggest that the error calibration for SESCA should be carried out using synthetic data, for which the errors in the reference data are known.

Finally, the error profiles of method V3 indicates that the estimated SS deviations in Section 3.1 of the SP175 set were indeed overestimated. SS deviations obtained by method V5 suggest an average 10% error for the SP175 reference structures. Further, estimating the model errors using different basis sets yield more consistent results with method V5 than that of V3, highlighting that V5 is more robust to the choice of the basis set.

In addition to the model validation accuracy, we also quantified how reliably model validation methods identify the correct SS composition, given a certain deviation from the ideal CD spectrum. To this aim, a ranking score *R_k_* for each synthetic spectrum *k* was determined using a given validation method. The ranking is given by the number of synthetic SS models with a lower or equal estimated error than the correct model of a synthetic data set. For our data set, the ranking for a spectrum can change between zero and twenty, with *R_k_*= 0 meaning, that the correct SS composition is uniquely identified by the validation method despite the errors in the CD spectrum.

In Figure 6C shows ranking scores of all five validation methods for representative synthetic spectra with non-SS contributions (C-subset). As the figure indicates, in the absence of CD errors all methods are able to identify the correct SS composition accurately, regardless of differences in their average accuracy. However, in the presence of 10% or larger non-SS contributions *R_k_* scores increase for all methods, indicating an increasing uncertainty of the true SS composition. This uncertainty is most likely due to the non-additive combination of CD and SS deviations, which entails partial error cancellation for certain model-spectrum combinations.

For a comparison, Fig. 6D depicts ranking scores for CD spectra with scaling errors only (S+ subset). The full effect of scaling errors is shown through method V1, for which the ranking scores increased similarly as seen for the non-SS contributions. Ranking scores for methods V2, V4, and V5, even in the presence of large scaling errors, remains zero due to re-scaling the CD spectra during validation. The effect of scaling errors is also reduced but not eliminated for V3, because here, the CD spectra are re-scaled based on the predicted spectrum of the (often incorrect) model SS composition. The SS deviations of the model combined with re-scaling and non-convergence of the deconvolution results in SS models within 6% deviation from the correct one showing the smallest apparent error, and therefore yielding a non-zero rank.

To compare the overall reliability of the five validation methods, the average and SD of the obtained ranking scores was also calculated over all synthetic spectra. As the values listed in Table 3 show, the average rank of V1 is close to 5, whereas V2 to V5 have similar average ranks between 2.8 and 3.0. The mean values and the large scatter of ranking scores between individual synthetic spectra suggest that, although V1 is less reliable for CD spectra with large scaling errors, the other four methods identify the correct SS composition with similar uncertainty.

Taken together, ranking scores and average model errors indicate that re-scaling the measured CD spectrum eliminates the effect of scaling errors and improves the reliability of model validation methods. However, non-SS contributions still impose an uncertainty on the estimated model errors and limit their precision. Calibration using synthetic CD data allowed us to take typical non-SS contributions into account and improve the accuracy of the SESCA model validation scheme compared to classical deconvolution-based methods, that neglect these contributions.

## 4 Conclusions

To interpret CD spectra of proteins in terms of estimating secondary structure content or validating putative model structures, several assumptions are required. These are accurately known reference secondary structures and protein concentrations during CD measurements, as well as negligible non-secondary structure contributions to the spectra. Using the SP175 reference set, we assessed and quantified to what extent these assumptions are fulfilled or violated. Our results suggest, that, even for the most accurate CD measurements, uncertainties in the protein concentration and non-SS contributions typically lead to 30% deviation of the measured spectrum from the true SS signal. In addition, typical reference SS compositions derived from X-ray crystallography or NMR spectroscopy also deviate from the SS composition during CD measurements by an average 10%, introducing further uncertainty to CD interpretation methods.

We also probed the effects of the observed CD and SS deviations on the accuracy of SS estimation, CD prediction, and model validation methods. To this aim, we constructed a synthetic reference data set of 21 CD spectra and SS compositions, for which we deliberately introduced known amounts of deviations based on those obtained for the SP175 set.

Testing the various methods on the synthetic data set shows that non-ideal CD spectra lead to errors in secondary structure estimation and decrease the accuracy of CD spectrum predictions. During the validation of structural models, typical CD deviations generally lead to the overestimation of the model error, and to a 5-15% uncertainty of the true SS composition. Although none of the tested model validation methods can eliminate the uncertainty, applying a method that takes the average CD deviations into account improves the model validation accuracy considerably. Our findings suggest that SESCA secondary structure estimation and model validation schemes can be improved based on the obtained distributions of CD deviations.

Using this new information, we implemented a new version of SESCA that automatically applies spectrum re-scaling during deconvolution and includes more accurate error parameters for model validation, obtained from systematic calibration based on synthetic CD data. These new results will also allow to go beyond determining the single SS composition that fits a given CD spectrum best and calculate the likelihood of all putative SS compositions for an improved uncertainty assessment.

## References

[1] B. M. Bulheller and J. D. Hirst. DichroCalc–circular and linear dichroism online. Bioinformatics, 25(4):539–540, February 2009.

[2] Gerald D. Fasman, editor. Circular Dichroism and the Conformational Analysis of Biomolecules. Springer US, Boston, MA, 1996.

[3] Fuchang Gao and Lixing Han. Implementing the Nelder-Mead simplex algorithm with adaptive parameters. Computational Optimization and Applications, 51(1):259–277, January 2012.

[4] Stanley C. Gill and Peter H. Von Hippel. Calculation of protein extinction coefficients from amino acid sequence data. Analytical biochemistry, 182(2):319–326, 1989.

[5] Sharon M. Kelly, Thomas J. Jess, and Nicholas C. Price. How to study proteins by circular dichroism. Biochimica et Biophysica Acta (BBA) – Proteins and Proteomics, 1751(2):119–139, August 2005.

[6] Daisuke Kihara. The effect of long-range interactions on the secondary structure formation of proteins. Protein Science, 14(8):1955–1963, August 2005.

[7] J. G. Lees, A. J. Miles, F. Wien, and B. A. Wallace. A reference database for circular dichroism spectroscopy covering fold and secondary structure space. Bioinformatics, 22(16):1955–1962, August 2006.

[8] A. Mahrenholz, T. T. Andersen, Y. Bao, S. A. Cohen, N. D. Denslow, J. Hulmes, P. E. Hunziker, K. Mann, K. M. Schegg, and K. West. ABRF Amino acid analysis survey: Identification of proteins electroblotted to PVDF. 1997.

[9] Parthasarathy Manavalan and W. Curtis Johnson. Protein secondary structure from circular dichroism spectra. Journal of Biosciences, 8(1-2):141–149, 1985.

[10] David Mao, E. Wachter, and B. A. Wallace. Folding of the mitochondrial proton adenosine triphosphatase proteolipid channel in phospholipid vesicles. Biochemistry, 21(20):4960–4968, September 1982.

[11] András Micsonai, Frank Wien, Linda Kernya, Young-Ho Lee, Yuji Goto, Matthieu Réfrégiers, and József Kardos. Accurate secondary structure prediction and fold recognition for circular dichroism spectroscopy. Proceedings of the National Academy of Sciences, 112(24):E3095–E3103, June 2015.

[12] Gabor Nagy, Maxim Igaev, Nykola C. Jones, Søren V. Hoffmann, and Helmut Grubmüller. SESCA : Predicting Circular Dichroism Spectra from Protein Molecular Structures. Journal of Chemical Theory and Computation, August 2019.

